## Supplementary Figures and materials for "Mecp2 Fine-tunes Quiescence Exit by Targeting Nuclear Receptors"

**Supplementary Figure Legends**

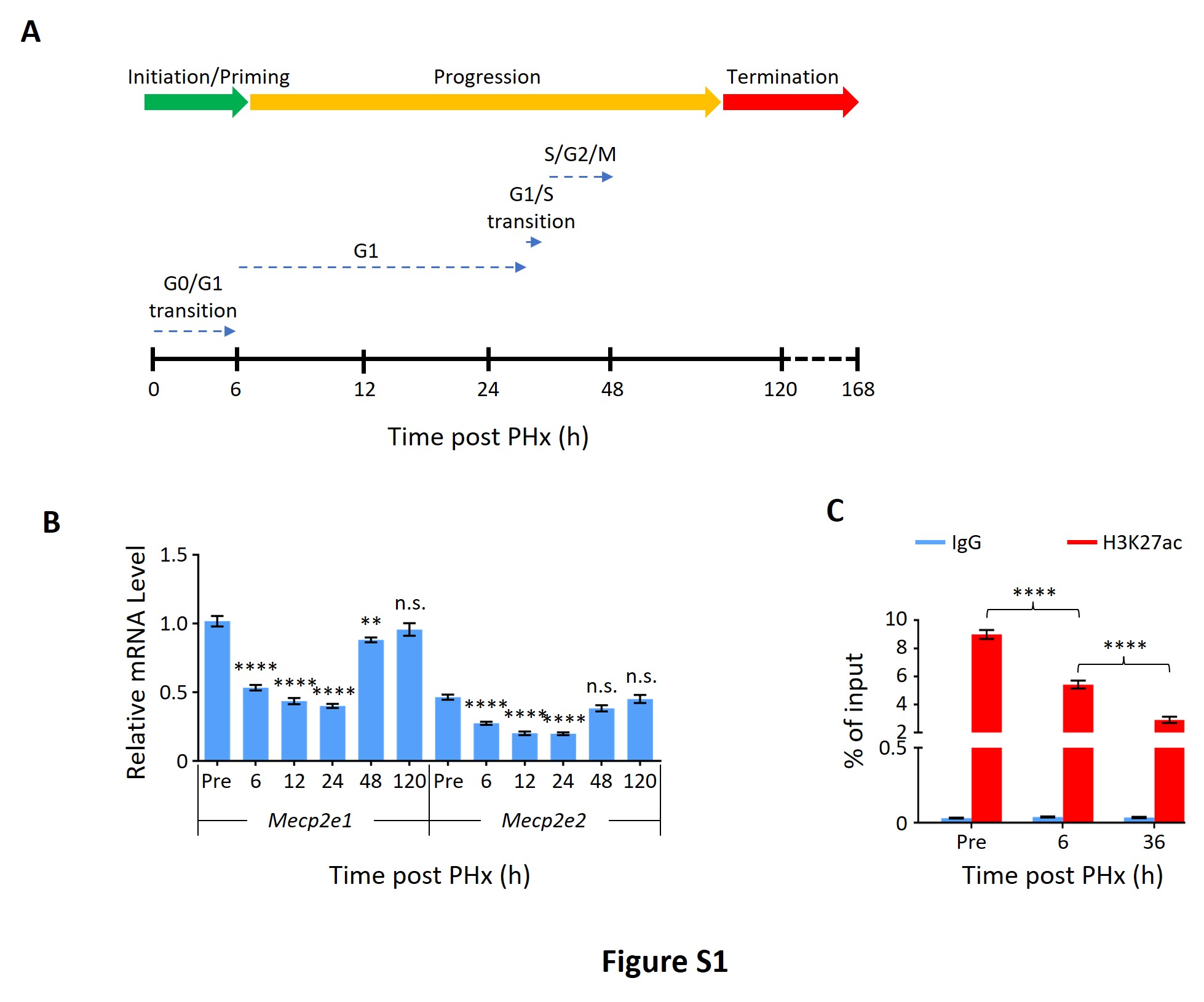

**Figure S1. Mecp2 is dynamically expressed during PHx-induced liver regeneration**

(A) Schematic for the PHx time course and data collection. Three phases of liver regeneration and the cell cycle progression of the first round of cell division after PHx are indicated by rainbow-colored and dotted arrows, respectively.
(B) Real-time PCR to evaluate mRNA levels of two isoforms of Mecp2, *Mecp2e1* and *Mecp2e2*, at different time points after PHx. Data are presented as means ± SEM; n = 6; n.s., not significant; ***p* < 0.01; *****p* < 0.0001 by one-way ANOVA.
(C) ChIP-qPCR showing H3K27ac occupancy at the *Mecp2* promoter at the indicated time points after PHx. IgG served as a negative control. Data are presented as means ± SEM; n = 6; *****p* < 0.0001 by two-way ANOVA.

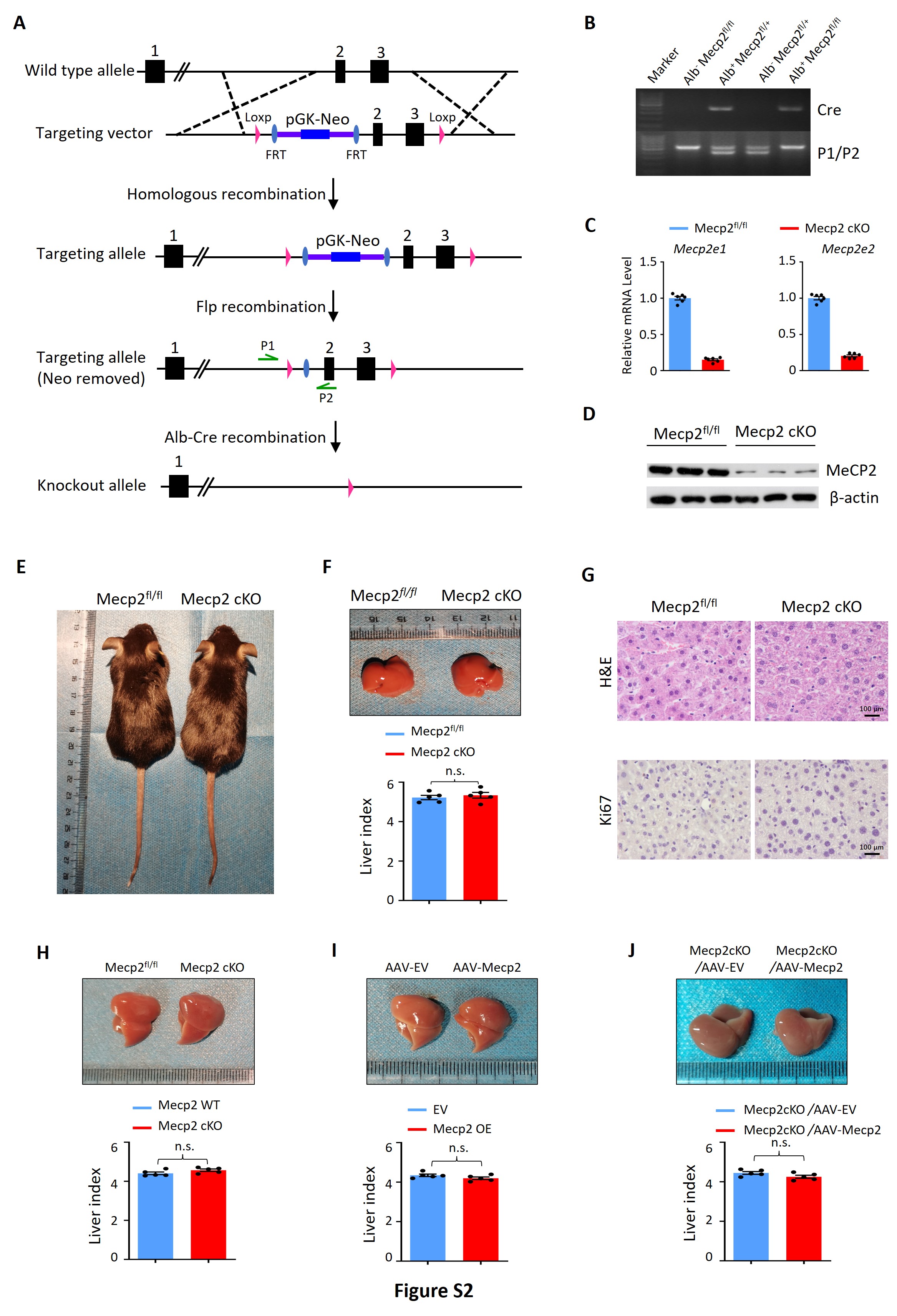

**Figure S2. Modulation of Mecp2 expression in the mouse liver.**

(A) Schematic diagram of loss of Mecp2 in mouse hepatocyte. In the Mecp2 floxed allele, Mecp2 exon 2 and 3 were flanked with loxp sites. The Neo-cassette was removed by mating with FLP recombinase homozygous mice. The exon 2 and 3 was deleted by mating with Alb-Cre recombinase homozygous mice.

(B-D) Evaluation of hepatocyte-specific depletion of Mecp2 by genotyping (B), real-time PCR (C), and WB (D) in Mecp2^fl/fl^ and Mecp2 cKO livers. Data are presented as means ± SEM. n = 6 mice/group; *****p* < 0.0001 by one-way ANOVA.

(E) Representative images of Mecp2^fl/fl^ and Mecp2 cKO mice at 3 months of age.
(F) Representative liver morphology and liver index of Mecp2^fl/fl^ and Mecp2 cKO mice before PHx. Data are presented as means ± SEM. n = 5 mice/group; n.s. not significant by Student’s t-test.
(G) Representative H&E staining and IHC for Ki67 images of liver tissues from Mecp2^fl/fl^ and Mecp2 cKO mice before PHx.
(H) Representative liver morphology and liver index of Mecp2^fl/fl^ and Mecp2 cKO mice at 7 days post-PHx. Data are presented as means ± SEM. n = 5 mice/group; n.s. not significant by Student’s t-test.
(I) Representative liver morphology and liver index of Mecp2^fl/fl^ mice without (AAV-EV) or with (AAV-Mecp2) AAV-mediated Mecp2 overexpression at 7 days post-PHx. EV, empty vector. Data are presented as means ± SEM. n = 5 mice/group; n.s. not significant by Student’s t-test.
(J) Representative liver morphology and liver index of Mecp2-cKO mice without (Mecp2 cKO/AAV-EV) or with (Mecp2 cKO/AAV-Mecp2) the rescue of Mecp2 at 7 days post-PHx. Data are presented as means ± SEM. n = 5 mice/group; n.s. not significant by Student’s t-test.

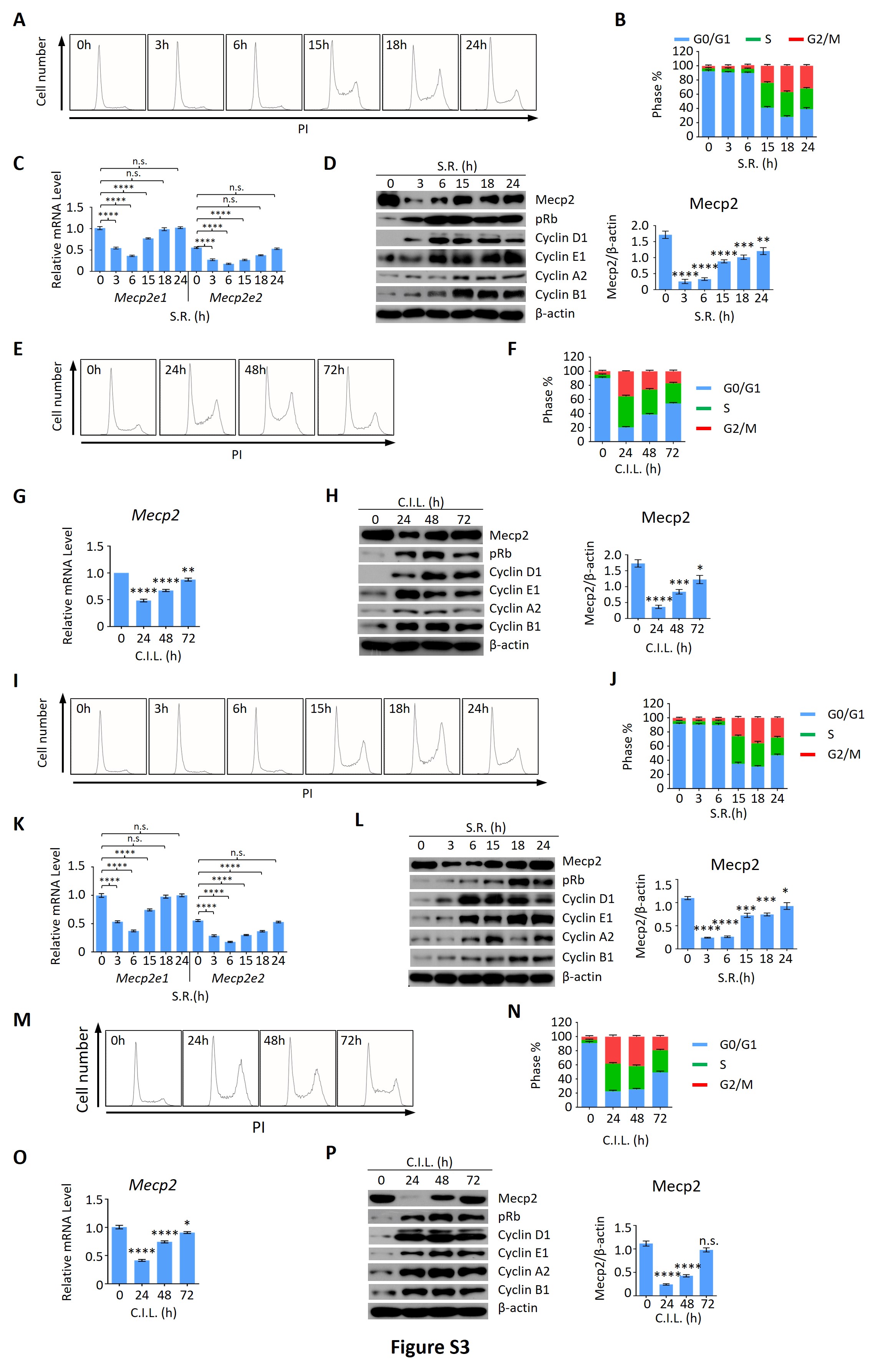

**Figure S3. Mecp2 is dynamically expressed during quiescence exit in both HT22 and HUVECs.**

S.R.-induced (A-D, and I-L) or C.I.L.-induced (E-H and M-P) quiescence exits in mouse hippocampal neuronal HT22 cells (A-H) or in HUVECs (I-P) was analyzed by PI staining (A, B, E, F, I, J, M and N), real-time PCR (C, G, K and O) and WB (D, H, L and P) in a time course of quiescence exit. (A, E, I, and M) Representative histograms of PI staining. (B, F, J, and N) Statistical analysis of the cell cycle distribution. (D, H, L, and P) WB of Mecp2, pRb, Cyclin D1, Cyclin E1, Cyclin A1, Cyclin B1 and β-actin in cells released from G0. Left panel, quantification of Mecp2 protein levels. In (C) and (K), n=9; In (O) and (P), n=6; In (D), (G), (H) and (L) n = 3; Data are presented as means ± SEM; n.s., not significant; **p* < 0.05; ***p* < 0.01; ****p* < 0.001; *****p* < 0.0001 by one-way ANOVA.

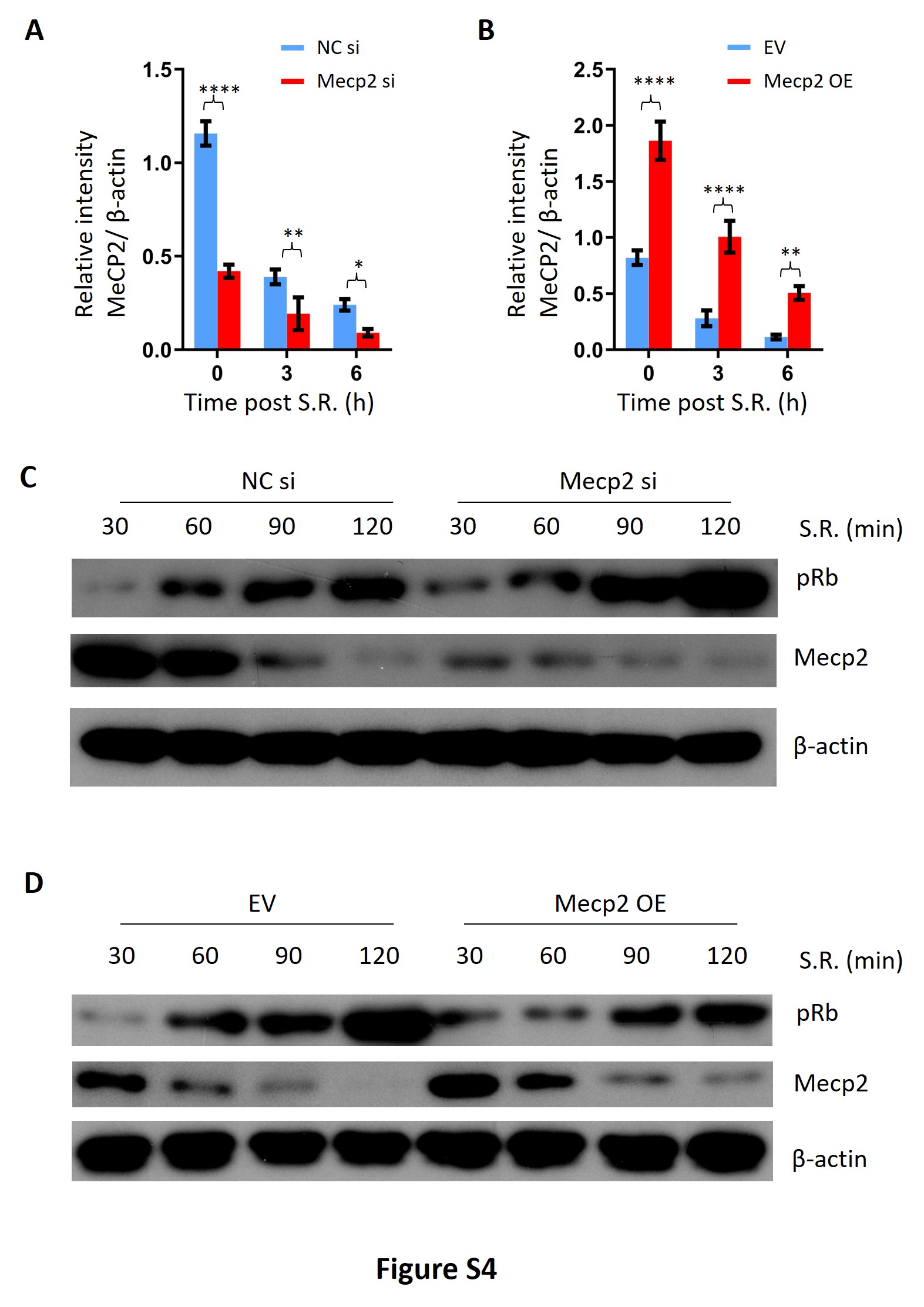

**Figure S4. Mecp2 negatively regulates the G0/G1 transition in the cellular model of S.R.-induced quiescence exit.**

(A) Protein quantitation of MeCP2 protein expression normalized to β-actin for Fig 4B Western Blotting. Data are presented as means ± SEM; n =3. **p* < 0.05; ****p* < 0.001; *****p* < 0.0001 by two-way ANOVA.

(B) Protein quantitation of MeCP2 protein expression normalized to β-actin for Fig 4F Western Blotting. Data are presented as means ± SEM; n =3. ***p* < 0.01; *****p* < 0.0001 by two-way ANOVA.

(C) WB of Mecp2 and pRb in control and Mecp2 KD 3T3 cells released from S.S.-induced quiescence at the indicated time points.

(D) WB of Mecp2 and pRb in control and Mecp2 OE 3T3 cells released from S.S.-induced quiescence at the indicated time points.

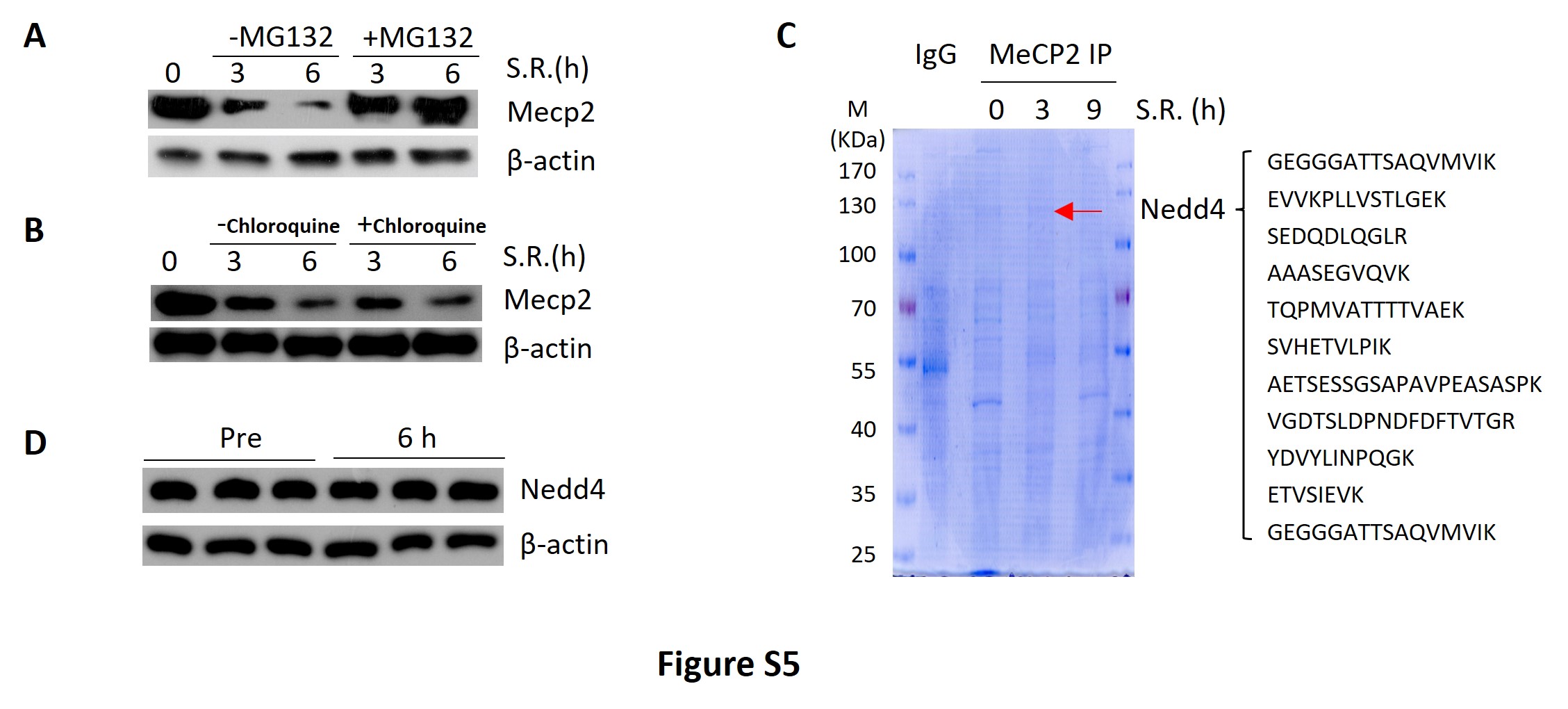

**Figure S5. Nedd4 interacts with Mecp2**

(A) WB of Mecp2 in quiescent 3T3 cells treated with or without MG132 after being released from G0 by S.R..

(B) WB of Mecp2 in quiescent 3T3 cells treated with or without chloroquine after being released from G0 by S.R..

(C) Coomassie staining of Mecp2 co-IP proteins separated by SDS-PAGE. Right, representative peptides of Nedd4 identified by mass spectrometry.

(D) WB showing protein levels of Nedd4 in mouse livers before and 6 h after PHx.

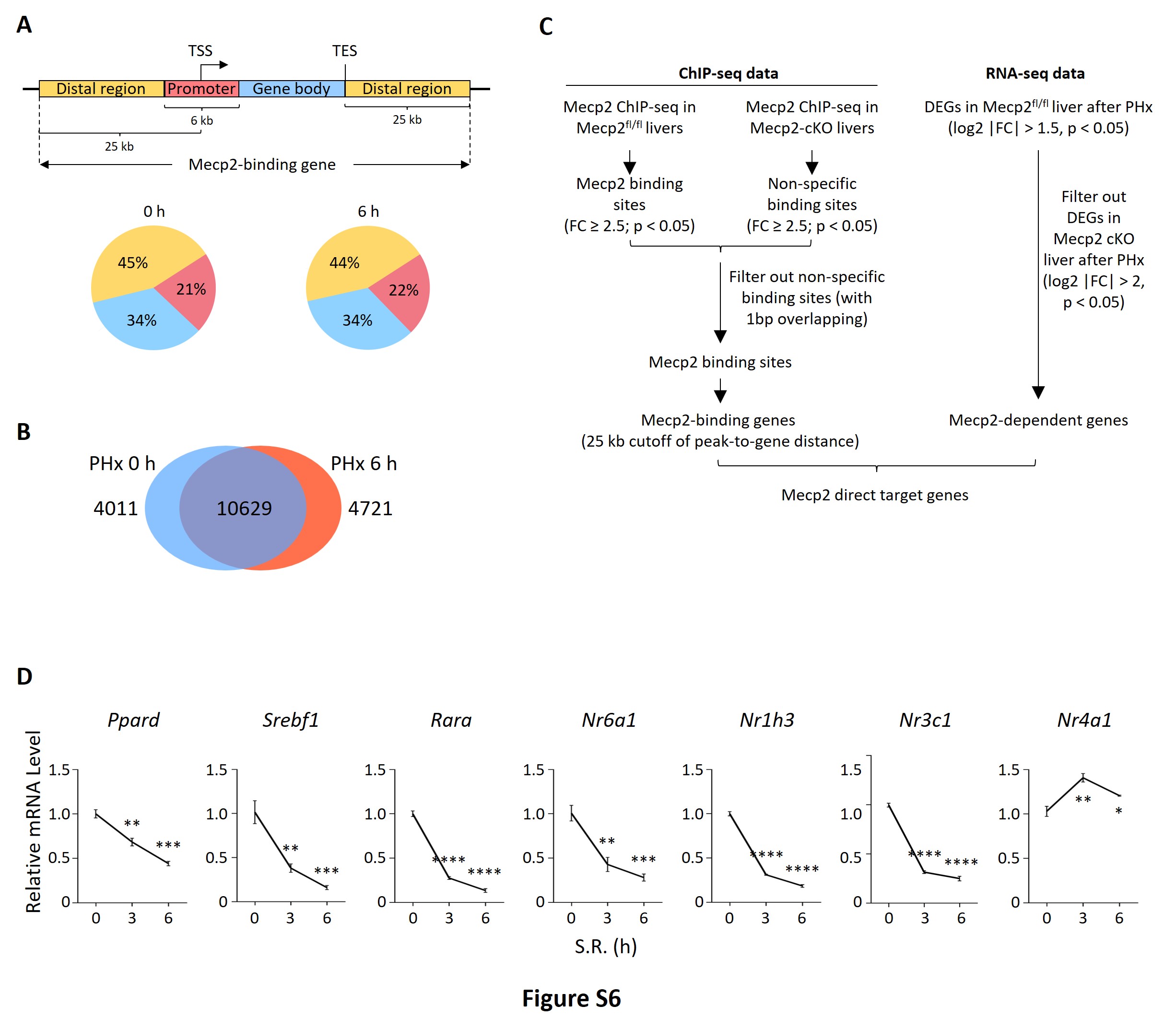

**Figure S6. Related to Figure 6.**

(A) Schematic showing the definition of Mecp2-binding genes. Lower panel, distribution of Mecp2 peaks in defined promoter-proximal, gene body and distal regions. TSS, transcriptional start site; TES, transcriptional end site. The cutoff of distance of peak-to-gene is 25 kb.
(B) Venn diagram overlap of Mecp2 binding genes in Mecp2^fl/fl^ livers before (0 h) and after PHx (6 h).
(C) Pipeline showing the integration of Mecp2 ChIP-seq data with RNA-seq data to identify Mecp2-direct target genes.
(D) Real-time PCR validation of PHx-repressed NRs in 3T3 cells upon S.R.-induced quiescence exit. Data are presented as means ± SEM; n = 3. **p* < 0.05; ***p* < 0.01; ****p* < 0.001; *****p* < 0.0001 by one-way ANOVA.

**Resources table**

| REAGENT or RESOURCE | SOURCE | IDENTIFIER |
| --- | --- | --- |
| Antibodies | | |
| Albumin | Proteintech | Cat#66051-1-Ig, RRID:AB_11042320 |
| β-actin | Proteintech | Cat#66009-1, RRID:AB_2687938 |
| Cyclin A2 | abcam | Cat# ab38, RRID:AB_304084 |
| Cyclin B1 | abcam | Cat# ab72, RRID:AB_305751 |
| Cyclin D1 | abcam | Cat#ab16663, RRID:AB_443423 |
| Cyclin E1 | Cell Signaling Technology | Cat#20808, RRID:AB_2783554 |
| H3K27ac | abcam | Cat#ab4729, RRID:AB_2118291 |
| IgG | Cell Signaling Technology | Cat#2729, RRID:AB_1031062 |
| K48-ubiquitin | abcam | Ca#ab140601 |
| Ki67-Immunofluorescence | abcam | Cat#ab279653 |
| Ki67-FACS | Biolegend | Cat#652406, RRID:AB_2561930 |
| MeCP2 | Cell Signaling Technology | Cat#3456, RRID:AB_2143849 |
| MeCP2-ChIP | abcam | Cat#ab2828, RRID:AB_2143853 |
| Nedd4 | Proteintech | Cat#21698-1-AP, RRID:AB_10858626 |
| Nr1h3 | Proteintech | Cat#14351-1-AP |
| p-Rb S807/811 | Cell Signaling Technology | Cat#8516, RRID:AB_11178658 |
| Rara | Proteintech | Cat#10331-1-AP |
| Ubiquitin | Cell Signaling Technology | Cat#3936, RRID:AB_331292 |
| Donkey anti-Mouse IgG (H+L) Highly Cross-Adsorbed Secondary Antibody, Alexa Fluor 488 | Thermo Fisher Scientific | Cat#A21202 RRID:AB_141607 |
| Donkey anti-Rabbit IgG (H+L) Highly Cross-Adsorbed Secondary Antibody, Alexa Fluor 594 | Thermo Fisher Scientific | Cat#A21207 RRID:AB_141637 |
| Bacterial and virus strains | | |
| pLKO.1 vector | Addgene | #8453 |
| pCMV-deltaR8 | Addgene | #12263 |
| pCMV-VsVg | Addgene | #8454 |
| AAV8- TBG-GFP | Genechem Co., Ltd | N/A |
| AAV8-TBG-MeCP2 | Genechem Co., Ltd | GOSV0233517 |
| Chemicals, peptides, and recombinant proteins | | |
| MG132 | selleck | Cat#S2619 |
| Puromycin | Sigma-Aldrich | Cat#P8833 |
| Polybrene | Sigma-Aldrich | Cat#TR-1003 |
| Pen Strep | Gibco | Cat#15140-122 |
| Complete Protease Inhibitor mini EASY packs EDTA-Free | Roche | Cat#05892791001 |
| Lipofectamine™ 3000 Transfection Reagent | Thermo Fisher Scientific | Cat#L3000-015 |
| Propidium Iodide (PI)/RNase Staining Solution | Cell Signaling Technology | Cat#4087 |
| DAPI | Thermo Fisher Scientific | Cat#D-1306 |
| Pierce™ IP Lysis Buffer | Thermo Fisher Scientific | Cat#87787 |
| Critical commercial assays | | |
| DAB Kit | ZSGB-BIO | Cat#ZLI-9018 |
| Western Lightning Plus ECL | Perkinelmer | Cat#0RT2655 |
| SimpleChIP® Plus Enzymatic Chromatin IP Kit | Cell Signaling Technology | Cat#9005 |
| Dynabeads™ Protein G | Thermo Fisher Scientific | Cat# 10004D |
| Coomassie Blue Super Fast Staining Solution | Beyotime | Cat# P0017F |
| Experimental models: Cell lines | | |
| NIH3T3 | ATCC | Cat#SCSP-515 |
| Ht22 | Procell Life Science&Technology | Cat#CL-0697 |
| HUVEC | ATCC | Cat#CRL-1730 |
| LentiX-293T | ATCC | Cat#ACS-4500 |
| Experimental models: Organisms/strains | | |
| B6. Cg-Speer6-ps1Tg (Alb-cre)21Mgn/J mice | The Jackson Laboratory | Cat# 003574 |
| MeCP2^flox/flox^ | Shanghai Model Organisms Center | Cat #NM-CKO-190001 |
| C57/BL6 | Guangdong Medical Laboratory Animal Center | N/A |
| Oligonucleotides | | |
| siRNA targeting sequence: MeCP2 #1: CCUGAAGGUUGGACACGAA | This paper | N/A |
| siRNA targeting sequence: MeCP2 #2: UGACAAAGCUUCCCGAUUA | This paper | N/A |
| siRNA targeting sequence: MeCP2 #3: CCGAAUUGCUGCUGCUUUA | This paper | N/A |
| siRNA targeting sequence: MeCP2 #4: CGAAAUGGCUGUGUAGCAA | This paper | N/A |
| siRNA targeting sequence: Nedd4 #1: CAGUGAUCCUUACGUAAGATT | This paper | N/A |
| siRNA targeting sequence: Nedd4 #2: GGGAAAUCGUACGAGAAGATT | This paper | N/A |
| siRNA targeting sequence: Nedd4 #3: GGAGGAUUAUGGGUGUGAATT | This paper | N/A |
| Primer for genotype: Alb, Forward:  ACCTGAAGATGTTCGCGATTATCT | This paper | N/A |
| Primer for genotype: Alb, Reserve:  ACCGTCAGTACGTGAGATATCTT | This paper | N/A |
| Primer for genotype: MeCP2, Forward:  GCTGGGGCCCTTGTTTTGAAT | This paper | N/A |
| Primer for genotype: MeCP2, Reserve:  GCTTTAGGTTGCTGGTGATA | This paper | N/A |
| Primer for qRT-PCR: Ar, Forward:  CTGGGAAGGGTCTACCCAC | This paper | N/A |
| Primer for qRT-PCR: Ar, Reserve:  GGTGCTATGTTAGCGGCCTC | This paper | N/A |
| Primer for qRT-PCR: β-actin mouse, Forward:  GGCTGTATTCCCCTCCATCG | This paper | N/A |
| Primer for qRT-PCR: β-actin mouse, Reserve:  CCAGTTGGTAACAATGCCATGT | This paper | N/A |
| Primer for qRT-PCR: β-actin human, Forward:  CACCATTGGCAATGAGCGGTTC | This paper | N/A |
| Primer for qRT-PCR: β-actin human, Reserve:  AGGTCTTTGCGGATGTCCACGT | This paper | N/A |
| Primer for qRT-PCR: MeCP2 common, Forward:  TATTTGATCAATCCCCAGGG | This paper | N/A |
| Primer for qRT-PCR: MeCP2 common, Reserve:  CTCCCTCTCCCAGTTACCGT | This paper | N/A |
| Primer for qRT-PCR: MeCP2 mouse, Forward:  GAGCGGCACTGGGAGACC | This paper | N/A |
| Primer for qRT-PCR: MeCP2 mouse, Reserve:  CTGGATGGTGGTGATGAT | This paper | N/A |
| Primer for qRT-PCR: MeCP2 human, Forward:  GATGTGTATTTGATCAATCCC | This paper | N/A |
| Primer for qRT-PCR: MeCP2 human, Reserve:  TTAGGGTCCAGGGATGTGTC | This paper | N/A |
| Primer for qRT-PCR: Nedd4, Forward:  TCGGAGGACGAGGTATGGG | This paper | N/A |
| Primer for qRT-PCR: Nedd4, Reserve:  GGTACGGATCAGCAGTGAACA | This paper | N/A |
| Primer for qRT-PCR: Nr1h3, Forward:  CTCAATGCCTGATGTTTCTCCT | This paper | N/A |
| Primer for qRT-PCR: Nr1h3, Reserve:  TCCAACCCTATCCCTAAAGCAA | This paper | N/A |
| Primer for qRT-PCR: Nr1i3, Forward:  ATATGGGCCGAGGAACTGTGT | This paper | N/A |
| Primer for qRT-PCR: Nr1i3, Reserve:  GGCGTGGAAATGATAGCCTGT | This paper | N/A |
| Primer for qRT-PCR: Nr2f6, Forward:  GAGGACGATTCGGCGTCAC | This paper | N/A |
| Primer for qRT-PCR: Nr2f6, Reserve:  GTAATGCTTTCCACTGGACTTGT | This paper | N/A |
| Primer for qRT-PCR: Nr3c1, Forward:  AGCTCCCCCTGGTAGAGAC | This paper | N/A |
| Primer for qRT-PCR: Nr3c1, Reserve:  GGTGAAGACGCAGAAACCTTG | This paper | N/A |
| Primer for qRT-PCR: Nr4a1, Forward:  TTGAGTTCGGCAAGCCTACC | This paper | N/A |
| Primer for qRT-PCR: Nr4a1, Reserve:  GTGTACCCGTCCATGAAGGTG | This paper | N/A |
| Primer for qRT-PCR: Nr5a2, Forward:  TGAGGAACAACTCCGGGAAAA | This paper | N/A |
| Primer for qRT-PCR: Nr5a2, Reserve:  CAGACACTTTATCGCCACACA | This paper | N/A |
| Primer for qRT-PCR: Nr6a1, Forward:  CGCAACGGTTTCTGTCAGGAT | This paper | N/A |
| Primer for qRT-PCR: Nr6a1, Reserve:  GTTCAGCTCGATCATCTGGGA | This paper | N/A |
| Primer for qRT-PCR: Ppard, Forward:  CTCATGAATGTGCCCCAGGT | This paper | N/A |
| Primer for qRT-PCR: Ppard, Reserve:  GTGCAGCAAGGTCTCACTCT | This paper | N/A |
| Primer for qRT-PCR: Rxrg, Forward:  CATGAGCCCTTCAGTAGCCTT | This paper | N/A |
| Primer for qRT-PCR: Rxrg, Reserve:  CGGAGAGCCAAGAGCATTGAG | This paper | N/A |
| Primer for qRT-PCR: Rara, Forward:  ATGTACGAGAGTGTGGAAGTCG | This paper | N/A |
| Primer for qRT-PCR: Rara, Reserve:  ACAGGCCCGGTTCTGGTTA | This paper | N/A |
| Primer for qRT-PCR: Srebf1, Forward:  GCAGCCACCATCTAGCCTG | This paper | N/A |
| Primer for qRT-PCR: Srebf1, Reserve:  CAGCAGTGAGTCTGCCTTGAT | This paper | N/A |
| Primer for ChIP-QPCR: MeCP2, Forward:  TAAGTGACAGGAGTCACAGCG | This paper | N/A |
| Primer for ChIP-QPCR: MeCP2, Reserve:  TGGGACGTTGTATGTAACGGG | This paper | N/A |
| Primer for ChIP-QPCR: Nr1h3, Forward:  CAGCACGTTGTAATGGAAGCC | This paper | N/A |
| Primer for ChIP-QPCR: Nr1h3, Reserve:  TAGCATTCAGTGGAGGGAAGG | This paper | N/A |
| Primer for ChIP-QPCR: Rara, Forward:  CGATGAGTGGCAAGGTCTTT | This paper | N/A |
| Primer for ChIP-QPCR: Rara, Reserve:  ATAGCATAGCACCAGGGACAC | This paper | N/A |
| shRNA for Nr1h3: CCTCAAGGACTTCAGTTACAA | This paper | N/A |
| shRNA for Rara: GAGCAGCAGTTCCGAAGAGAT | This paper | N/A |
| Recombinant DNA | | |
| Mecp2 Mouse Tagged ORF Clone, transcript variant 1 | OriGene | Cat#MR226839 |
| Mecp2 Mouse Tagged ORF Clone, transcript variant 2 | OriGene | Cat#MR207745 |
| Nedd4 Mouse Tagged ORF Clone | OriGene | Cat#MR222243 |
| pCMV6-Entry Mammalian Expression Vector | OriGene | Cat# PS100001 |
| Software and algorithms | | |
| Image J | Image J | https://imagej.nih.gov/ij/ |
| FlowJo 10 | BD Biosciences | https://www.flowjo.com/ |
| GraphPad Prism 5.0 | Prism | http://www.graphpad-prism.cn/prism.html |
| ModFit 3.3 | ModFit LT | https://www.vsh.com/products/mflt/index.asp |
